# The gut microbiome promotes detoxification responses to an environmental toxicant

**DOI:** 10.1101/2025.08.14.670327

**Authors:** Ian N. Krout, Rie Matsuzaki, Alexandria C. White, Sherry Tsui, Lisa Blackmer-Raynolds, Sean D. Kelly, Jianjun Chang, Mattie Braselton, Priya E. D’Souza, Catherine E. Mullins, Parinya Panuwet, Volha Yakimavets, Dana B. Barr, Douglas I. Walker, W. Michael Caudle, Timothy R. Sampson

## Abstract

At the host-environment interface, the indigenous microbiome is poised to facilitate interactions with exogenous components. Here, we show that the microbiome is necessary for metabolic and transcriptional detoxification responses to the neurotoxic pyrethroid insecticide, deltamethrin. While oral deltamethrin exposure shapes gut microbiome composition, it is not directly microbially metabolized. Instead, we observe microbiome-dependence on host hepatic and intestinal detoxification responses, with diminished activity in germ-free mice. Colonization with a complex microbiome in adulthood maintained limited hepatic responses, suggesting developmental contributions. However, mono-colonization with specific microbes increased colonic expression of a key detoxification enzyme, revealing a protective role for active microbial signaling in the colon. Overall, our data demonstrate that the microbiome is necessary to prime and activate a host response against a model environmental toxicant. Through both developmental and active signaling across organ compartments, these data support that the microbiome may contribute to risk and outcomes of toxicant-associated disease.

**Highlights:**

- The gut microbiome mediates the host response to environmental toxicants.
- Key xenobiotic metabolism genes are modulated by the microbiome
- Early life signaling is necessary to promote hepatic responsiveness to toxicants in adulthood.
- Specific and active microbial signaling promotes colonic detoxification gene expression.

## INTRODUCTION

The native gut microbiome encodes for >50-fold more genes than their mammalian hosts, highlighting their genetic capacity for myriad physiological functions that can impact health and disease (Cani, 2018; Cirstea et al., 2023; Zhu et al., 2010). Activities within or derived from the gut microbiome are essential for digestion and nutrient acquisition, immune system maturation and function, brain development and behavior, among other central roles (Dekaboruah et al., 2020; Sommer and Backhed, 2013). In line with their contributions to dietary metabolism, specific, native gut bacterial species have been demonstrated to directly degrade pharmaceuticals and other xenobiotics (Cirstea et al., 2023; Culp et al., 2024; Dekaboruah et al., 2020; Garcia-Santamarina et al., 2024; van Kessel et al., 2019; Zimmermann et al., 2019a). Through these actions, direct microbial metabolism can impact pharmacokinetics and ultimately, drug efficacy (Cirstea et al., 2023; Collins and Patterson, 2020; Feng et al., 2020; Garcia-Santamarina et al., 2024; Li et al., 2016; Maini Rekdal et al., 2019; van Kessel et al., 2019; Weersma et al., 2020). While direct metabolism by the gut microbiome is one action on xenobiotics, signals derived from the gut microbiome may further stimulate or prime physiological systems that act to metabolize exogenous molecules (Cai et al., 2022; Feng et al., 2020; Koppel et al., 2017). Such actions of indigenous microbes on environmental toxicants, such as prevalent chemical pollutants including pesticides, are largely undescribed.

Pyrethroid pesticides are one class of environmental contaminants that are highly prevalent due to widespread residential, commercial, and agricultural use (Saillenfait et al., 2015). While an essential tool to combat vector-borne diseases and the treatment of parasitic arthropod infections (Ravula and Yenugu, 2021), this family of pesticide has emerged as an exposure concern due to its abundance in foodstuffs (Saillenfait et al., 2015). Pyrethroids are neurotoxins and exert their effects in part through prolonged activation of voltage-gated sodium channels and inhibition of GABA_A_ receptors (Vijverberg and van den Bercken, 1990). Epidemiological and experimental evidence suggests associations between pyrethroid exposure and specific neurological diseases including Parkinson’s disease and attention-deficit and hyperactivity disorders (Elwan et al., 2006; Kannarkat et al., 2015; Narayan et al., 2017; Richardson et al., 2015; Ritz et al., 2016; Wagner-Schuman et al., 2015; Wang et al., 2011). Given these disease associations, as well as a primarily oral route of exposure through food, it is essential to understand how the local intestinal microbiome is impacted by exposure and whether the microbiome has a contribution to the metabolism of this toxicant.

Here, we use the pyrethroid pesticide, deltamethrin, to directly test contributions by the microbiome to host detoxification responses. In a mouse model of oral exposure, we found that an intact microbiome is necessary for the metabolic and detoxification response of deltamethrin. While exposure selected for distinct microbiome compositions both *in vivo* and *ex vivo*, we did not observe deltamethrin metabolism directly by the microbiome. Rather, we found that the indigenous microbiome promoted xenobiotic responses in both the hepatic and intestinal systems. The hepatic compartment was strikingly non-responsive in the absence of a microbiome. Colonization with a complex microbiome in adulthood maintained limited responsiveness to toxicant exposure in the hepatic compartment, indicative of developmental impacts. However, we found that specific microbes actively modulated colonic expression of *Ces2a,* a key xenobiotic detoxification enzyme indicating a specific and active role for the microbiome in protecting the colon. Overall, these data demonstrate a central role for the native microbiome in maintaining a physiological state that is capable of responding appropriately to environmental exposures. Understanding specific microbes and their metabolites which give rise to these effects will reveal an intrinsic microbial contributor to the variability of outcomes across environmental exposures.

## RESULTS

### Oral deltamethrin impacts the composition of the gut microbiome

In order to first determine the effects of deltamethrin exposure on the gut microbiome, we used a mouse model of chronic, oral exposure. Wildtype mice were treated once weekly for 12wks by oral gavage with 3mg/kg deltamethrin (a dose which falls within established no observable effect and lowest observable adverse effect limits) or with a corn oil vehicle (White et al., 2025). Assessment of stool microbiome composition by 16S RNA sequencing over this 12wk period revealed a significant compositional shift (Figure 1). We observed a small, yet significant increase in alpha diversity, as assessed by the Simpson index, in mice of either sex orally exposed to deltamethrin (Figures 1A, 1B, 1F, and 1G). In addition, beta diversity measurement by the Bray-Curtis metric demonstrated a substantial shift in community structure of those mice exposed to deltamethrin, with limited differences in vehicle exposed animals (Figures 1C and 1H). While the microbiomes of both male and female mice were impacted by exposure, they differed in the specific bacterial taxa affected (Figures 1D and 1I, and Supplemental Table S1). For instance, in male mice we observe an increase in *Akkermansia sp*. when vehicle-treated and *Allobaculum sp*. in the deltamethrin-treated group (Figure 1E), which do not appear in female mice. Female animals, however, do display an increase in *Turicibacter sp*. to a greater extent in mice exposed with deltamethrin (Figure 1J). Both sexes displayed a significant increase in *Bifidobacterium sp.* (Figures 1E and 1J, and Supplemental Table S1) upon oral exposure to deltamethrin, but with differing temporal patterns. Overall, chronic ingestion of a low concentration of insecticide leads to a significant shift in the composition of the gut microbiome.

**Figure 1.**
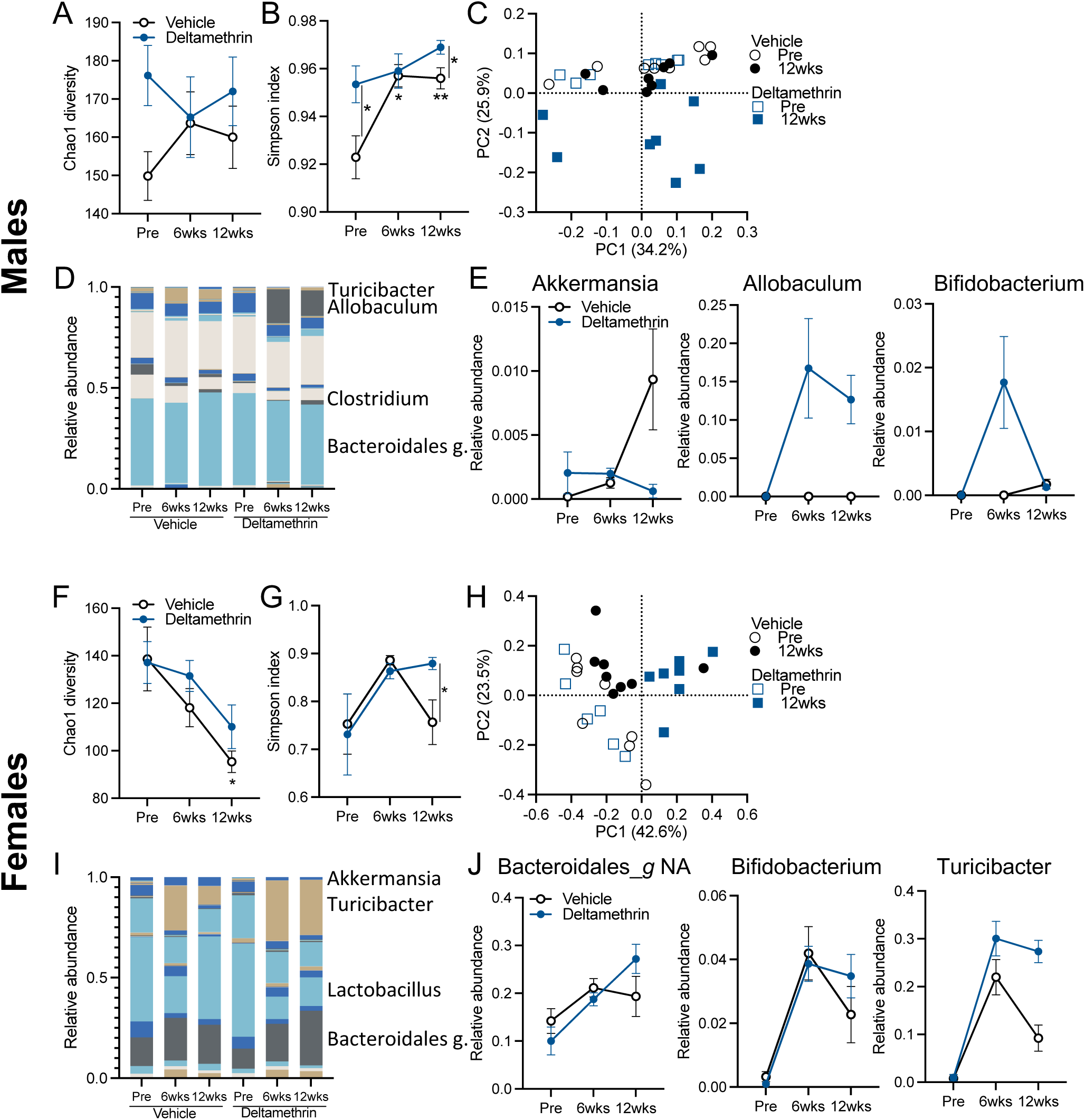
Chronic deltamethrin ingestion sex-specifically impacts the composition of the gut microbiome. Male (**A-E**) and female (**F-J**) wildtype mice were orally exposed with 3mg/kg deltamethrin or corn oil vehicle, 1x weekly for 12 weeks. Prior to exposure, and at 6 and 12wks, stool was subjected to 16S microbiome profiling. (**A, B, F, G**) Alpha diversity measurements for Chao1 (**A, F**) and Simpson indexes (**B, G**). (**C, H**) PCA plots of Bray-Curtis beta diversity measures. (**D, I**) Genera composition charts. (**E, J**) Select genera displaying time or treatment effects (see Supplemental Table S1). N=8; **A, B, E,-G, J** points represent mean and bars the standard error. **C, H** points represent individuals. **A, B, F, G** data assessed by mixed-effects REML and post-hoc Fisher’s tests, with lines indicating between group differences. \**p*≤0.05; \*\**p* ≤0.01

Oral deltamethrin exposure results in an acute constipation phenotype (White et al., 2025), which may be responsible for the compositional shift in the microbiome, rather than direct effects of the toxicant on bacterial physiology. To test how deltamethrin impacted the microbiome composition directly, we performed *ex vivo* exposures using murine stool-derived cultures. Anaerobic culture of murine stool in the presence of deltamethrin, or the related pyrethroids bioallethrin and permethrin, at two concentrations resulted in a significant loss of diversity over 72hrs, albeit less than vehicle-treated cultures (Figures S1A and S1B). Overall, the community structure which arose in culture following deltamethrin treatment differed from vehicle controls (Figure S1C). Surprisingly, despite established toxicodynamic differences, each pyrethroid resulted in a similar community shift, suggesting shared, direct effects on bacterial growth or survival (Figures S1A - S1D). While we did not observe any genera statistically different from the vehicle treated group, a number of genera were qualitatively impacted. Opposite from the *in vivo* exposures, genera including *Allobaculum, Bifidobacterium* and *Lactobacillus* were qualitatively decreased (Figures S1D and S1E). In line with this, we performed *in vitro* growth assessments of specific *Bifidobacterium* and *Lactobacillus* species to identify whether pyrethroids inhibited their growth. We found that treatment of these mono-cultures with deltamethrin resulted in *Bifidobacterium* growing slightly accelerated at low doses, but no species tested was fully inhibited in its growth, even at the highest doses (Figure S1F). Indeed, little difference in growth of *Lactobacillus* was observed at any dose (Figure S1F). This may suggest that some species are able to utilize deltamethrin metabolically or are resistant to its effects. Overall, while deltamethrin can directly affect the composition of the microbiome, its effects on the community *in vivo* are likely not due to direct action of the toxicant on bacterial targets and instead may be due to the effects of the toxicant on the host and intestinal environment.

### A complex microbiome is necessary for the detoxification of the pyrethroid insecticide, deltamethrin

Given that we observed both direct and indirect effects *in vivo* and *ex vivo* on microbiome composition upon deltamethrin treatment, we next tested whether the microbiome contributed to the detoxification and metabolism of deltamethrin. We performed oral exposures in germ-free wild-type mice in comparison to mice harboring a conventional microbiome, and measured the concentrations of 3-phenoxybenzoic acid (3-PBA, deltamethrin’s first-order metabolite in its detoxification). At 4hrs post-exposure (C_MAX_), we observe substantially decreased 3-PBA in serum, with qualitative decrease in matched urine, in germ-free mice compared to conventionally-colonized controls (Figure 2A and 2B). However, the ratio of 3-PBA in urine vs serum was identical irrespective of colonization status, suggesting that excretion of the metabolite is similar between groups (Figure 2C). Thus, the presence of an intact, native microbiome is necessary for the efficient *in vivo* metabolism of deltamethrin to 3-PBA. In order to test whether the microbiome was directly metabolizing deltamethrin, as has been observed for some pharmaceutical compounds (Garcia-Santamarina et al., 2024; Zimmermann et al., 2019a, b), we measured the loss of deltamethrin in *ex vivo* murine-derived stool microbiome cultures. Even at 48hrs post-exposure, we were unable to detect a significant loss of deltamethrin, relative to media-only controls (Figure 2D). Therefore, while the presence of a native microbiome is necessary for the host to metabolize deltamethrin, this effect is not due to microbial action on the toxicant directly.

**Figure 2.**
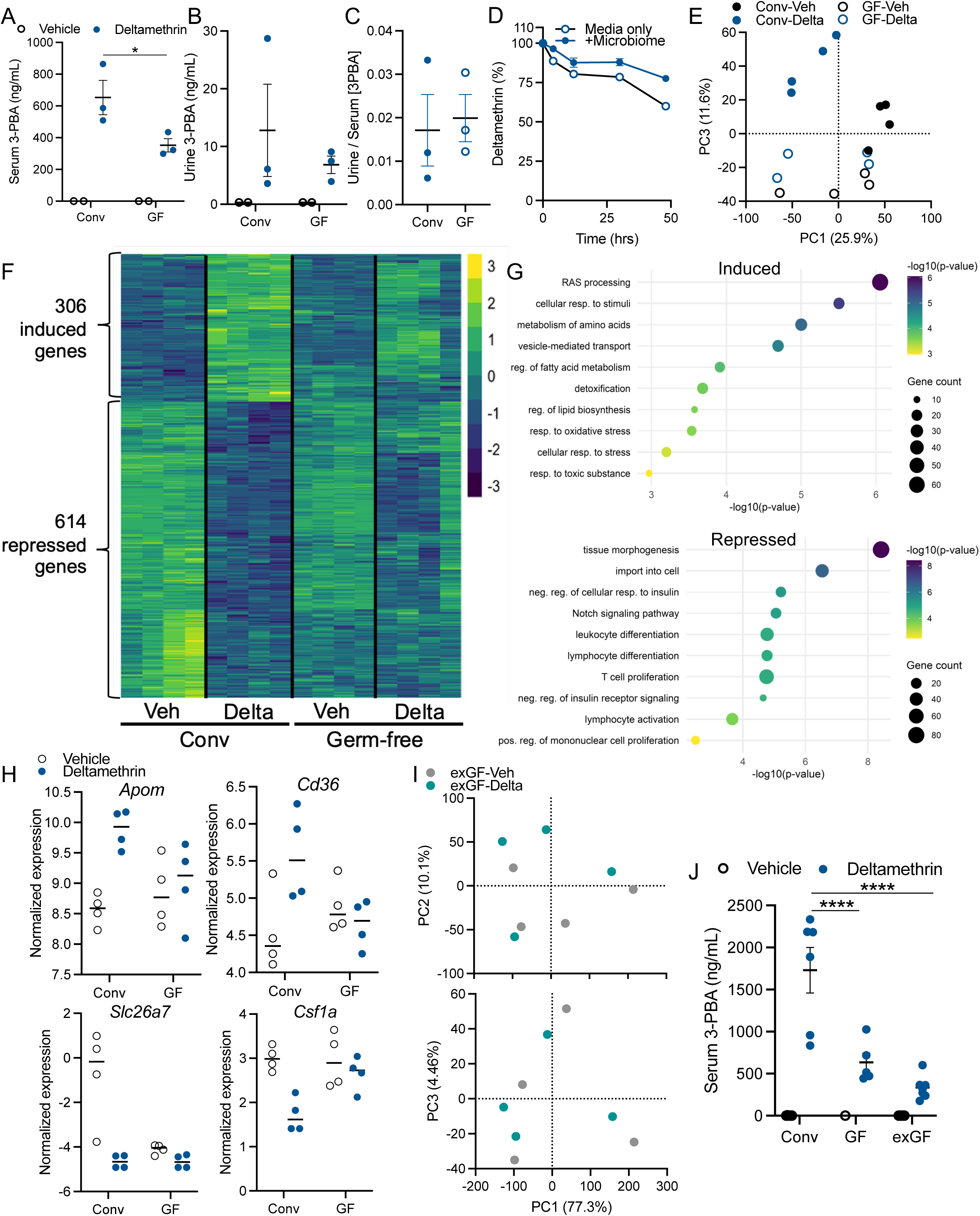
A native microbiome is developmentally necessary for deltamethrin metabolism and hepatic toxicant responses *in vivo*. Concentrations of 3-phenoxybenzoic acid (3-PBA) in serum (**A**) and urine (**B**) of wildtype conventional and germ-free mice at 4hrs post oral exposure with 3mg/kg deltamethrin. **C**) Urine/serum excretion ratio. **D**) Percent loss of deltamethrin in *ex vivo* anaerobic stool cultures. **E-H)** RNAseq analysis of liver tissue derived from conventional and germ-free mice. **E)** PCA comparing transcriptomes induced by deltamethrin between conventional and GF mice. **F)** Heatmap of 920 DEGs identified in conventional mice, and their relative expression in GF mice. **G)** Metascape pathway analysis of enriched pathway terms for genes increased and decreased in conventional mice treated with deltamethrin. **H)** Select DEGs, isolated from data in **F**, exemplifying lack of responsiveness in GF mice. **I)** PCA comparing RNAseq of hepatic tissues of exGF mice following deltamethrin exposure. **J**) Serum 3-PBA concentrations in serum of conventional, germ-free, and ex-GF mice. N=2-3, **A-C;** N=4, **E-I;** N=1-6, **J**, **A-C, E, H-J**, points/columns represent individuals, and bars the mean and standard error; **D** points represent the mean and bars the standard error. **A-D**, **J** Data assessed by two-way ANOVA and post-hoc Fisher’s tests, with lines indicating between group differences. DEGs determined by FC>[1] and adjusted *p-*value <0.05. \**p*≤0.05, \*\*\*\**p* ≤0.0001

### Irreversible signals derived from the gut microbiome promote host hepatic detoxification responses

Since the native microbiome did not directly metabolize deltamethrin *in vitro*, we sought to address how the presence of the indigenous microbiome contributes to host-mediated toxicant metabolism. We first performed transcriptomic analysis of hepatic tissue, as the site of first-pass metabolism of xenobiotics. In line with prior reports (Selwyn et al., 2015), we found that an indigenous microbiome is necessary for the steady-state hepatic transcriptional landscape. Bulk RNAseq analysis revealed that livers of exposure-naïve, wildtype, germ-free mice displayed a subtle, yet significant transcriptional shift compared to conventional animals (Figures S2, S3 and Supplemental Table S2). Specifically, we found 16 differentially expressed genes (DEGs; fold-change ≥[1], adjust. *p*-value<0.05, 9 increased, 7 decreased), dependent on the presence of a microbiome (Figure S2A). Pathway analysis of those pathways increased in GF mice (repressed by microbial signals) revealed an enrichment in genes involved in metabolism-namely precursor synthesis and small molecule biosynthesis (Figure S2B and Supplemental Table S2), while no enriched pathways amongst repressed genes were detected in GF-derived liver tissue. While many of the DEGs are involved in xenobiotic responses, we most notably observed that GF mice had a significant decrease in the expression of *Cyp3a11*, a cytochrome P450 enzyme (CYP) involved in a number of detoxification pathways (Figure S2C) and known to be induced by microbiome-derived bile acids (Goodwin et al., 2003). Thus, demonstrating that microbial signals are necessary for maintaining homeostatic expression of xenobiotic detoxification genes.

Given this observation of microbiome-dependent hepatic state, we next sought to determine whether microbiome-derived signals are necessary for the transcriptional responsiveness to deltamethrin. At 4hrs post-exposure to oral deltamethrin or vehicle control, hepatic tissues from colonized and GF mice were subjected to RNAseq analysis. We observed that deltamethrin evoked a robust, acute transcriptional response in livers of conventionally-colonized mice (Figure 2E and Supplemental Table S3). A total of 920 DEGs (306 increased, 614 decreased) were elicited in hepatic tissue following deltamethrin exposure (Figure 2F). Notably, we observed transcriptional responsiveness of numerous CYP genes including increased *Cyp2c29*, *Cyp2c70*, *Cyp4a32*, *Cyp2b10*, involved in various levels of xenobiotic metabolism (Figure S3 and Supplemental Table S3). Pathway analysis revealed an increase in detoxification pathways (*e.g.* GO 0098754), as expected, as well as pathways involved in oxidative stress responses (GO 0006979) and amino acid metabolism (R-MMU-71291) (Figure 2G and Supplemental Table S3). Hepatic pathways that were decreased following deltamethrin exposure included insulin signaling (*e.g.* GO 1900077), in line with our prior report in enteroendocrine cells (White et al., 2025), as well as a downregulation of numerous immune and inflammatory pathways (*e.g.* GO 0002521) (Figure 2G and Supplemental Table S3), similar to systemic observations on immune function post-pyrethroid exposure (Suwanchaichinda et al., 2005).

In striking contrast, livers derived from GF mice exposed to deltamethrin displayed no DEGs, and PCA indicated no overall shift in the transcriptomes of GF mice following deltamethrin exposure (Figure 2E). Thus, an intact microbiome is necessary for a robust hepatic transcriptional response to this pyrethroid toxicant. Comparison of those deltamethrin-modulated DEGs in conventional mice to their responses in the absence of a microbiome emphasizes the necessity of microbiome-derived signaling for host responsiveness to toxicant exposure (Figure 2F). This lack of responsiveness is observed broadly and is represented in select genes involved in key transcriptional pathways, including increased pathways in detoxification and oxidative stress (GO 0098754, *e.g. Apom, Cd36*) and decreased pathways such as immune function (*e.g.* GO 0002521, *e.g. Csf1a*) (Figure 2H). We therefore conclude that the native microbiome is necessary to promote the host hepatic response to deltamethrin, allowing a detoxification and protective response.

The indigenous microbiome impacts host physiology through active, constant signaling that may be modulated in adulthood, and/or via developmental processes that are engrained in early life (Sharon et al., 2016; Sommer and Backhed, 2013). To begin to differentiate these processes, we colonized GF mice in adulthood with wildtype, exposure-naïve, mouse microbiomes (*i.e.* exGF mice). Following a 4hr oral deltamethrin exposure, as performed in our prior experiments, we analyzed hepatic transcriptomes by RNAseq. Nearly identical to our findings in GF mice, we found that livers derived from exGF mice remained transcriptionally non-responsive to oral deltamethrin exposure with only one DEG detected (Supplemental Table S4). PCA of the hepatic transcriptome highlighted no substantial shift in the overall gene expression (Figure 2I), demonstrating that the non-responsive phenotype in the absence of a microbiome remains through adulthood. Supporting this persistent dampened hepatic response to deltamethrin, exGF mice maintained a decreased ability to metabolize deltamethrin to 3-PBA (Figure 2J). Similar to GF mice, exGF mice showed significantly lower serum 3-PBA concentrations at 4hrs post-oral exposure. We therefore conclude that the hepatic response to toxicant exposure must be entrained through early-life or developmental signals by the microbiome and is not actively modulated or reversable by microbiome colonization in adulthood.

### Intestinal pyrethroid toxicity is actively modulated by the indigenous microbiome

Given the intimate association between the gastrointestinal tract and the microbiome, we sought to determine whether the microbiome contributions to hepatic toxicant responses were mirrored within colonic tissues. First, we assessed the baseline, microbiome-dependent transcriptional landscape between conventionally-colonized and GF mice. In line with prior reports (Fu et al., 2017), we observed a robust, microbiome-dependent colonic transcriptome (Figure S4A and Supplemental Table S5). We found 247 microbiome-dependent DEGs, with 69 increased and 178 decreased in GF mice compared to colonized controls. As expected, these mapped to a variety of established microbiome-dependent pathways including regulation of hormone metabolism (*e.g.* GO 0010817) and immune function (*e.g.* GO 0002252). In these colonic tissues, we also observed significant, microbiome-dependent regulation of genes involved in various xenobiotic responses and detoxification pathways (*e.g.* GO 0009410, GO 0098754, and GO 0042537) (Figure S4 and Supplemental Table S5). Specifically, we found a number of genes for CYP enzymes dysregulated in the absence of microbiome signals. This includes significant decreases in the expression of *Cyp3a11*, *Cyp3a44*, *Cyp3a25*, *Cyp2c55*, and increased *Cyp2d34*, *Cyp2d10*, *Cyp2d13*, *Cyp2d9*, *Cyp2f2*, *Cyp1a1* (Figure 3A). Some of these key enzymes contribute to metabolism of microbiome-derived bile acids and other lipids (*e.g. Cyp3a*, *Cyp2d9*)(Goodwin et al., 2003), however many do not suggesting that microbial influences extend beyond canonical endogenous pathways, to include enzymes more commonly recognized for their role in xenobiotic detoxification. Taken together, these findings further implicate a critical role for microbiome-derived signals in maintaining broader steady-state levels of detoxification machinery for pyrethroids and other xenobiotics within colonic tissue.

**Figure 3.**
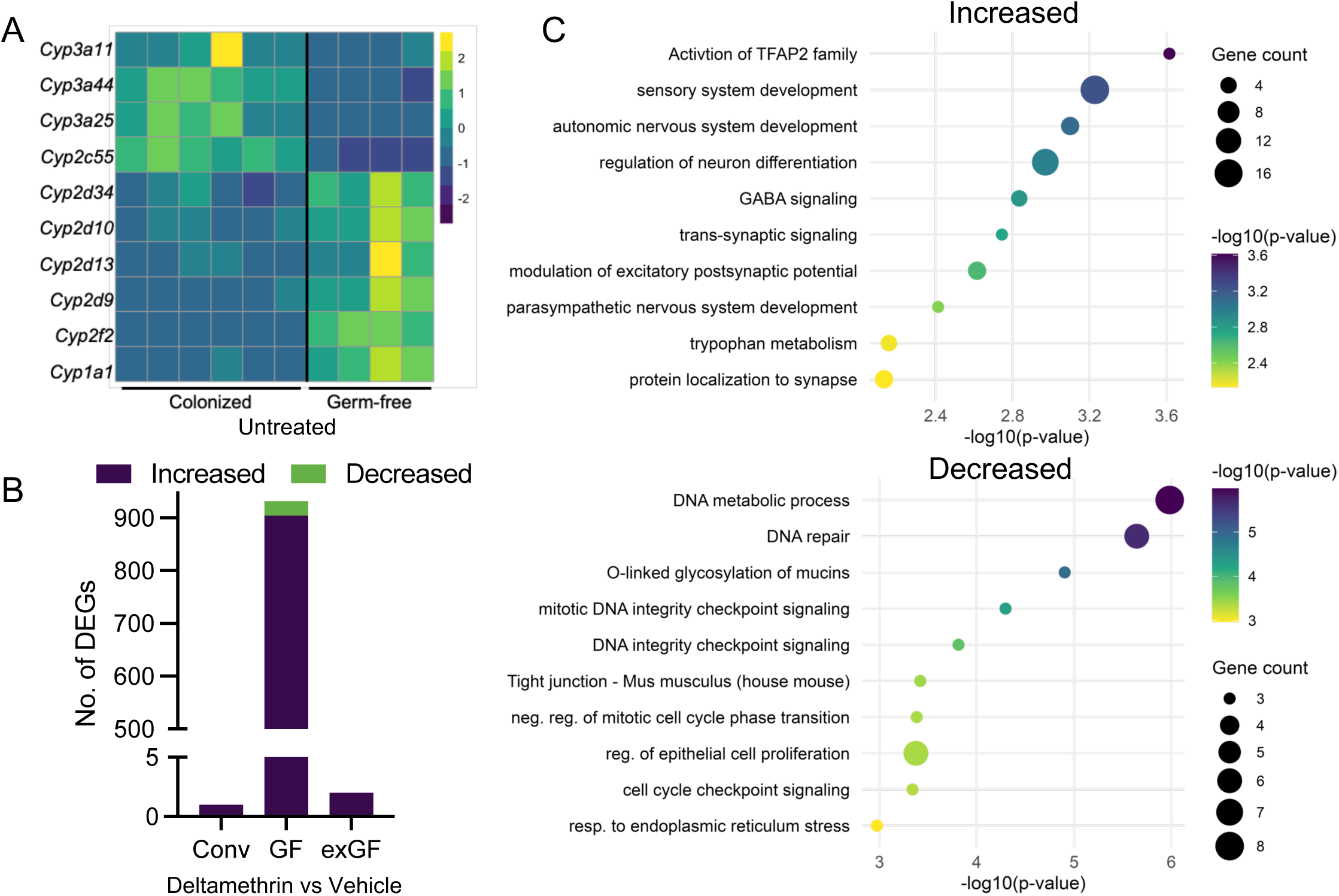
The gut microbiome actively limits colonic toxicity responses to deltamethrin exposure . **A**) Differentially expressed CYP genes within colonic tissue derived from germ-free mice in comparison to colonized controls. **B)** Number of differentially expressed genes in colonic tissue following deltamethrin exposure, based on colonization status. **C)** Metascape pathway analysis of enriched pathway terms for genes increased or decreased in germ-free mice colons following exposure to deltamethrin. N=4, 6, columns in **A** represent individual mice. DEGs determined by FC>[1] and adjusted *p-*value <0.05.

Following a 4hr oral deltamethrin exposure, as performed in our prior experiments, we analyzed colonic transcriptomes by RNAseq to identify both toxicant-induced and microbiome-dependent responses, as we found in the hepatic compartment. Unlike our observations in the liver, we found that colonic tissue derived from mice with an intact microbiome did not undergo a strong transcriptional response. In fact, only one gene, *GM3248*, was significantly upregulated following deltamethrin exposure (Figure 3B and Supplemental Table S6). Surprisingly, colons from GF mice underwent a robust transcriptional response with 932 DEGs (904 upregulated, 28 downregulated; Figure 3B and Supplemental Table S7). In contrast to the hepatic response in colonized mice, the genes induced in the colon of GF mice following deltamethrin exposure were not involved in detoxification responses (Figure 3C and Supplemental Table S7). Rather, deltamethrin-induced genes in colonic tissue of GF mice were characterized by roles in synaptic activity (*e.g.* GO 0099550), driven by increased expression of critical synaptic receptors including *Adra1a, Adrb2, Grin2a, Grin2b*, *Gabra2,* and *Gabrab1*. These are notable since many are directly targeted by pyrethroids (*e.g.*, GABA_A_ receptors [*Gabra2, Gabrab1*]) or modulated by activation of pyrethroid-sensitive voltage-gated sodium channels and thought to be responsible for pyrethroid toxicity (Birk et al., 2015; Gammon and Casida, 1983; George et al., 2012; Rinne et al., 2013; Soderlund et al., 2002), In addition, we observed substantial enrichment in genes involved in neuronal development (*e.g.* GO 0048483) due to increased expression of central regulatory genes including *Tfap2a, Tfap2b, Tfap2c*, *Sox11*, and *Hoxb1*,*2*, further implicating the microbiome in protection against pyrethroid-induced toxicity through regulation of these cell fate pathways. Enrichment of genes repressed in GF mice following deltamethrin exposure were largely characterized by pathways in DNA repair and cell cycle checkpoint signaling (*e.g.* GO 0006281, GO 0044774, and GO 0000075). This is despite no significant alteration of the known pyrethroid targets, voltage-gated sodium channels and GABA_A_ receptors, in the absence of an intact microbiome (Supplemental Table S5). Together, these strongly suggest a microbiome-dependent impact on local pyrethroid toxicity in the colon.

To investigate whether the microbiome-dependent intestinal response was due to developmental signals (as we observed in the hepatic system), or active microbial signaling, we assessed the colonic transcriptome of GF mice colonized in adulthood with a complex microbiome. As expected, these exGF mice showed a substantially shifted baseline colonic transcriptome from both GF and conventionally colonized mice (Figure S4B and Supplemental Table S8). We found 1352 DEGs, with 1151 increased and 201 decreased in exGF mice compared to colons derived from conventionally-colonized animals (Supplemental Table S8). Despite this significant effect on the baseline colonic transcriptome, deltamethrin exposure of exGF mice showed little transcriptional responsiveness in the colons (with only two DEGs: *Ighv1-5* and *Prss27*), recapitulating the lack of colonic response in colonized mice (Supplemental Table S6). This suggests that while the microbiome has a developmental impact on hepatic detoxification functions for pyrethroids, the microbiome is necessary for preventing local pyrethroid toxicity in the colon through active, reversable signaling.

### The native microbiome actively modulates intestinal detoxification enzymes

To begin to explore how active, microbiome-derived signals may alter toxicological outcomes in the intestine, we examined whether expression of microbiome-dependent CYP genes (Figure 3A) were reversable following colonization in adulthood. We found that colonization fully reversed the expression of dysregulated CYPs from GF mice (Figure 4A and Supplemental Table S9), back to levels observed in normally-colonized mice. Thus, the expression of microbiome-dependent, colon-resident CYP genes are actively modulated by the microbiome. Pyrethroid toxicants are acted on by specific carboxylesterases including the *Ces* family present in both hepatic and intestinal tissue. We did not detect any *Ces* gene differentially regulated in the hepatic compartment by either the microbiome (Supplemental Table S2) or by deltamethrin (Supplemental Table S3). Within the colon, however, the pyrethroid hydrolase *Ces2a* was significantly repressed in the absence of an intact microbiome (Supplemental Table S5 and Figure 4B). In the GI tract, *Ces2a* is primarily expressed within enterocytes of the epithelial layer poising it be acted on by both exogenous environmental chemicals and signals derived by the microbiome. Notably, we found that while Ces2a did not transcriptionally respond to deltamethrin exposure in either the liver or colon (Supplemental Tables S3, S6, and S7), it actively responded to colonization in adulthood with a complex microbiome (Figure 4B). In order to understand whether *Ces2a* expression is modulated by specific microbes, we mono-colonized GF mice in adulthood with a subset of representative type strains of native gut bacteria. We observed that only one organism in our panel, *Clostridum celatum*, resulted in *Ces2a* expression that was indistinguishable from wholly colonized mice (Figure 4C). Thus, specific microbial factors during colonization promote *Ces2a* expression rather than a general response to bacterial colonization. We interpret these data in total to suggest that *Ces2a*, and other key xenobiotic metabolizing genes, are actively and specifically modulated by the microbiome in their local intestinal tissue expression, in contrast to developmentally entrained pathways in the hepatic system.

**Figure 4.**
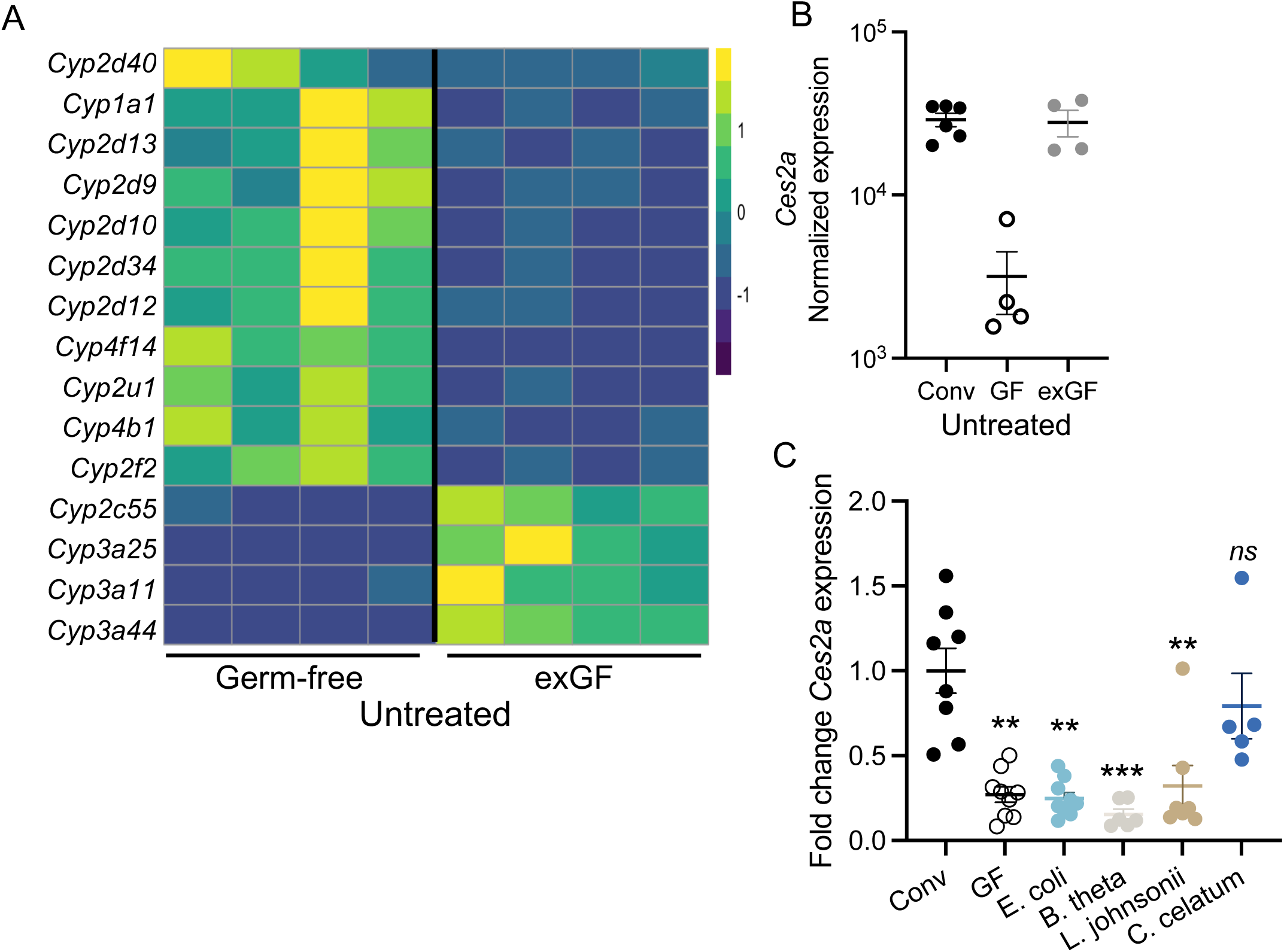
Microbiome-derived signals actively modulate intestinal xenobiotic response genes. **A)** Differentially expressed CYP genes within colonic tissue derived from germ-free mice in comparison to exGF mice. **B)** Normalized *Ces2a* expression by colonization status. **C)** qPCR for *Ces2a* in colonic tissue of mice monocolonized with the indicated bacterial species, in comparison to germ-free and conventional controls. N=4, 6, (**A, B**); N=5-9 (**C**) points/columns represent individuals, bars the mean and SEM. DEGs determined by FC>[1] and adjusted *p-*value <0.05. Data in **C** assessed by Kruskal-Wallis with post-hoc Dunn’s multiple comparisons test, compared to conventional controls.

## DISCUSSION

The native microbiota exerts a profound influence over all aspects of host organisms’ physiology through dynamic interactions with other organ systems across the lifespan (Cryan et al., 2019; Sharon et al., 2016; Sommer and Backhed, 2013). Situated at the physical interface between the host and the external environment, the gut microbiota is poised to facilitate interactions between exogenous stimuli and the host, such as modulating the biotransformation and systemic distribution of diverse exogenous compounds (Culp et al., 2024; Zimmermann et al., 2019a, b). These compounds include not only dietary chemicals and microbial factors, but also environmental toxicants such as pesticides and other xenobiotics (Koppel et al., 2017; Weersma et al., 2020). The capacity of the microbiota to both respond to and transform these chemical exposures, whether it be directly or indirectly, adds a layer of complexity to host detoxification pathways, with significant implications for metabolic, neurologic, and inflammatory disease risk (Chi et al., 2021; Singh et al., 2022). As such, exploring how microbiota is both impacted and facilitated by interactions with these chemicals is therefore critical to understanding the etiology and pathological heterogeneity of complex diseases.

With an estimated 10-fold the genetic capacity of the human genome (Ley et al., 2006; Qin et al., 2010), the bacterial component of the intestinal microbiome has an immense genetic repertoire by which to act on xenobiotics. The resident microbial community has been demonstrated to metabolize a wide range of physiologically important chemicals, including pharmaceutical compounds *in vitro* and *in vivo* (Culp et al., 2024; Garcia-Santamarina et al., 2024; Zimmermann et al., 2019a, b). Microbial actions, either by individual organisms or cross-species metabolism can modify pharmacological outcomes. For instance, specific microbes can act on levodopa, a primary therapeutic for Parkinson’s disease (PD) leading to loss of efficacy in model systems (Cirstea et al., 2023; Maini Rekdal et al., 2019; van Kessel et al., 2019). In addition, a robust set of data demonstrate precise contributions of individual microbes to the degradation of pharmaceutical compounds, altering pharmacokinetics (Zimmermann et al., 2019a, b). Separately, environmental microbiologists have identified a number of microbes with the ability to degrade ecologically-relevant xenobiotics in soil and water, as potential contributors to pollutant remediation (Masotti et al., 2023). Despite these microbial actions in the environment, contributions of the indigenous gut microbiome to the degradation of pesticide and other non-pharmaceutical toxicants during host exposures have been largely undescribed.

Our hepatic transcriptomic and metabolomic data revealed that microbial colonization was essential to induce a robust xenobiotic response to deltamethrin, and that this capacity could not be restored by introducing a complex community of microbes later in life. The murine liver undergoes significant post-natal maturation, when microbiota colonization occurs in parallel (Liang et al., 2022; Sharon et al., 2016). In our study, colonization of germ-free mice in early adulthood (8–10 wks of age) failed to restore hepatic responsiveness to deltamethrin, suggesting that microbial cues are required during a critical developmental window to establish normal xenobiotic responses in adulthood. This critical period may occur postnatally, prior to or during weaning; or even prenatally through maternal microbial signaling. Despite the specificity of the critical window, our findings support the existence of a microbiota-dependent period during which hepatic detoxification capacity is programmed. Once this window closes, early life microbiome-derived signaling results in long-lasting and irreversible contributions to xenobiotic responses. These results underscore the potential importance of early-life microbial exposures—such as those influenced by birth mode or breastfeeding—in shaping long-term susceptibility to toxicant-induced diseases.

In both hepatic and colonic tissues, we demonstrate that the microbiome is necessary for the homeostatic expression of a number of xenobiotic response genes. The microbiome-dependence in exposure-naïve mice is consistent with prior studies focused on hepatic physiology, demonstrating a microbiota-dependent influence on bile acid and other lipid metabolism (Togao et al., 2021; Zhang et al., 2014). One key to disentangling these findings lies in distinguishing which gene expression changes are driven by microbiota-mediated signaling versus those more directly responsive to xenobiotic exposures. Unlike in the hepatic compartment, colonic tissue exhibited enhanced responsiveness to deltamethrin in the absence of microbiota. Dysregulation of genes involved in synaptic function, neuronal activation, DNA repair, and cell cycle checkpoints suggest that the microbiota are protective against local colonic toxicity. In contrast to the irreversible metabolic deficits observed in the liver, this exaggerated colonic response was reversible in adulthood, suggesting active microbial signalling mediates local protection. Further, we demonstrated microbial-specificity in this process. The expression of *Ces2a*, a pyrethroid-hydrolyzing carboxylesterase (Stok et al., 2004), was microbiome-dependent, and mono-colonization with *Clostridium celatum*, but not other organisms tested, restored its expression. This supports the concept that both active and specific microbiome-derived signals modulate local host detoxification pathways. It is therefore plausible that microbiome composition at the time of exposure, alongside its contributions during development, contributes to the inherent variability of toxicological outcomes observed following an exposure event. One’s microbiome composition should therefore be considered as a potential risk factor in its interactions with a xenobiotic exposure. More coercively, our data raise the possibility that active manipulation of the microbiome, for instance with specific probiotic-type supplementation, may limit toxicity by promoting beneficial host responses. Our data show microbial specificity to this action, but do not identify the molecular drivers of this signaling pathway. Given the recognized metabolic versatility of *Clostridium* species and the production of bioactive metabolites by *C. celatum* (Guo et al., 2020), it remains to be clarified how *C. celatum* drives this effect and whether strain-specific interactions may differ substantially from community-level dynamics.

Pyrethroid insecticides, including deltamethrin, are environmentally prevalent and epidemiologically linked to a number of chronic diseases including metabolic disorders, neurological conditions, neurodegenerative diseases, and emerging links to cancer (Ascherio et al., 2006; Freire and Koifman, 2012; Narayan et al., 2017; Navarrete-Meneses and Perez-Vera, 2019; Paul et al., 2023; Ritz et al., 2016; Wagner-Schuman et al., 2015; Wang et al., 2011; Xie et al., 2024). Our data presented herein further support these associations. We observe that oral exposure to deltamethrin resulted in significant modulation of insulin response and lipid metabolism in the liver, in line with our prior report and highlighting impacts to metabolism that may shape metabolic syndrome (White et al., 2025). In susceptible GF mice, we see significant repression of cell cycle checkpoint genes with roles in limiting carcinogenesis, which may begin to provide insight to emerging human data linking pyrethroids to cancer (Navarrete-Meneses and Perez-Vera, 2019; Stok et al., 2004). In addition, we observe substantial dysregulation of genes involved in synaptic stability. Alterations of synaptic function in the intestinal tract and periphery are features of prodromal and early stages PD (Horsager et al., 2020; Horsager and Borghammer, 2024). Intriguingly, we and others observe that spore-forming bacteria, such as those closely related to *C. celatum*, are diminished in people living with PD (Romano et al., 2021; Wallen et al., 2022). Our data suggest that certain microbiome compositions, including those with diminished *C. celatum*, may exacerbate impacts of pyrethroid or other toxicant exposures. While we do not yet know the cause of the PD or other neurological disease-associated altered gut microbiome compositions, it is tempting to speculate that toxicant exposures may be one such factor which shifts the community. In line with human data and other animal models of pyrethroid exposure (Alhasson et al., 2017; Angoa-Perez et al., 2020; Zheng et al., 2024), we observe that deltamethrin increases PD-associated taxa including *Bifidobacterium*, *Akkermansia*, and *Turicibacter*, implicating shared pathological features within the intestinal environment between PD and toxicant exposure.

Together, our findings establish a critical role for the gut microbiota in shaping systemic and tissue-specific responses to environmental toxicants. While it is intuitive that distinct organs deploy context-specific programs to manage chemical stressors, our data reveal that these programs are differentially dependent on microbial input. In the hepatic system, microbiota-derived signals are essential to entrain a durable detoxification response in adulthood; whereas in the colon, active and specific microbial signals confer protective flexibility by modulating local expression of xenobiotic-processing genes. These contrasting dynamics underscore the importance of microbiota in programming organ-specific toxicological outcomes and raise fundamental questions about how microbe–host interactions are temporally and spatially integrated. Ultimately, our data support that early life microbial exposures and microbiome composition should be considered in environmental health risk assessments. Microbiota-targeted strategies may further offer promising avenues to protect the host organism from the diverse impacts of environmental toxicants.

## RESOURCE AVAILABILITY

### Lead contact

Further information and requests for resources should be directed to and will be fulfilled by the lead contact, Timothy Sampson.

### Materials availability

This study did not generate new, unique materials.

### Data and code availability

Transcriptomic and microbiome data are deposited at NIH SRA database, and source data and analysis at Zenodo, freely available upon acceptance.

Any additional information required to reanalyze the data reported in this paper is available from the lead contact upon request.

## ACKNOWLEDGMENTS

We thank Isabel Fraccaroli for scientific support; and all the members of the Emory Division of Animal Resources for technical support. We acknowledge support from the Emory Multiplexed Immunoassay Core (EMIC), the Emory Gnotobiotic Animal Core (EGAC) which are subsidized by the Emory University School of Medicine as Integrated Core Facilities and are supported by the Georgia Clinical and Translational Science Alliance of the NIH (UL1TR002378). This work is funded by NIH/NIEHS R01ES032440, Parkinson’s Foundation PF-JFA-830658, and an Emory HERCULES Pilot NIH/NIEHS P30ES019776 to TRS; NIH/NIEHS T32ES012870 to INK and ACW; and NIH/NIA F31AG076332 to LBR. The content is solely the responsibility of the authors and does not necessarily reflect the official views of the sponsors.

## AUTHOR CONTRIBUTIONS

TRS and WMC conceived and designed the study. TRS analyzed data and wrote the manuscript. INK, RM, ACW, ST, and LBR performed experiments. SDK and JJ performed animal manipulations, husbandry, and technical support. PP, VY, PD, and DBB performed in vivo metabolite measurements. CEM and DIW performed ex vivo metabolite measurements. All authors revised and approved the manuscript.

## DECLARATION OF INTERESTS

The authors have no conflicts of interest to declare.

## DECLARATION OF GENERATIVE AI AND AI-ASSISTED TECHNOLOGIES IN THE WRITING PROCESS

ChatGPT4 or other AI-assisted technologies were not used in the writing of this manuscript.

## STAR+METHODS

### EXPERIMENTAL MODEL AND SUBJECT DETAILS

#### Animal husbandry

Conventionally-reared C57Bl/6J (Jackson Laboratory, RRID: IMSR_JAX: 000664) male and female mice at eleven-to twelve weeks of age were housed (2-3 per cage) according to treatment and sex. Mice were provided water and standard, sterile mouse chow ad lib (LabDiet: 5001), and housed on a 12 h light/dark cycle. Germ-free (GF) C57Bl/6J mice were originally obtained via embryonic rederivation and bred within the Emory Gnotobiotic Animal Core for at least 3 generations prior to this study. GF mice were housed under microbiologically sterile conditions, within rigid ParkBio isolators. Culture and qPCR testing was performed on all autoclaved materials entering isolators (including food, water, and bedding) as well as monthly within the isolators themselves to confirm absence of microbial contamination. Generation of ex-GF mice were performed by 100uL oral gavage of wildtype, murine fecal slurries pooled from three, 11-week old, conventional C57Bl/6J and homogenized in 50 % glycerol (v/v) and 5 % sodium bicarbonate (w/v) into GF C57Bl/6J mice. GF and exGF status were confirmed via anaerobic culture on tryptic soy blood agar. Generation of mono-colonized mice was performed as described by our group previously(Blackmer-Raynolds et al., 2025; Hamilton et al., 2024). Briefly, type strains obtained from ATCC (as indicated in the Key Resource Table) were cultured under anaerobic conditions (5% H_2_, 10%CO_2_) in appropriate media. Overnight cultures were centrifuged, washed with PBS, and resuspended in 50 % glycerol (v/v) and 5 % sodium bicarbonate (w/v), and 100µL aliquots of ∼10^8^-10^9^ cfu were orally gavaged once. Mice remained in sterile housing, and upon the experimental endpoint, mono-colonization confirmed by culture on selective and non-selective medias.

At the experimental endpoints, 4hrs following final exposures, mice were humanely euthanized under deep isoflurane anesthesia via cardiac perfusion, for exsanguination and perfusion with sterile PBS. All biological tissues and matrices were immediately collected and flash frozen in liquid nitrogen. Samples were stored at –80 °C until further use. All animal husbandry and experimental procedures were performed in accordance with Emory’s Institutional Animal Care and Use Committee (IACUC) protocol #201900030.

## METHOD DETAILS

### Oral deltamethrin exposures

Mice received a 50 µL oral gavage of 3 mg/kg Deltamethrin (Chem Service, Cat#: N-11579-250MG) dissolved in filter-sterilized corn oil (Mazola) or a 50 µl gavage of filter-sterilized corn oil alone as a vehicle using a sterilized metal feeding needle (22G)(White et al., 2025). For acute exposures and measurement of transcriptional responses, a single exposure was performed 4hrs prior to humane euthanasia and tissue collection. For microbiome compositional assessments, mice were orally exposed 3x weekly for the duration of the experiment (12wks). Post-gavage all mice were monitored for signs of acute toxicity and sickness behaviors for the duration of the study.

### *Ex vivo* culture of stool microbiomes

Fecal pellets were sterilely collected from ten-to twelve-week-old, wildtype C57Bl/6J mice. A single pellet from each mouse (3 total mice) was immediately gently dissociated in 2 ml of sterile Brain Heart Infusion (BHI; BD Difco #237500) media. For microbiome compositional assessment, samples were briefly spun at 500 x *g* for 30sec and supernatants aliquoted into BHI containing deltamethrin (Chem Service, Cat#: N-11579-250MG), permethrin (Sigma, Cat#: 45614-250MG), or bioallethrin (Sigma, Cat#: 31489-250MG) at concentrations of 70µM or 140µM, as indicated. Treated samples were cultured for 72hrs at 37C under anerobic conditions (5% H_2_, 10%CO_2_), prior to collection for 16S sequencing.

For assessment of deltamethrin metabolism, supernatants from each mouse were placed into 1mM deltamethrin in BHI, and cultured at 37°C under anaerobic conditions. Aliquots were collected immediately, and at 4, 12, 24, and 48hrs, and compared to media alone, in the absence of added stool-derived microbes, and stored at -80°C prior to analysis.

For bacterial growth assays, frozen bacterial stocks were initially cultured anaerobically on solid medium (as indicated in the Key Resource Table) at 37°C for 24hrs. Single colonies were picked, and cultured overnight in liquid medium, and then diluted to standard OD600 for each species (0.05, *Bifidobacterium* and 0.10, *Lactobacillus*). Deltamethrin was diluted to 200µM in appropriate media, and serial dilutions performed in a 96-well plate. 1:10 dilutions of the indicated bacterial cultures were added, to a final volume of 300µl. Plate cultures were incubated anaerobically at 37°C, and OD600 was assessed every 2 hours. Growth occurred until the maximum optical density was reached (19hrs for *Bifidobacterium* and 12hrs for *Lactobacillus*). The percentage of max growth was determined based on the difference between the OD600 at each time point and the starting OD600 value used at time of inoculation.

### 16S microbiome compositional analysis

For *in vivo* studies, stool was sterilely collected prior to exposure, and at 6 and 12wks post exposure. For *ex vivo* studies, stool cultures were collected prior to and at 72hrs post-exposure. Samples were then subjected to 16S sequencing and analysis as we have performed previously(Hamilton et al., 2024) via full service 16S sequencing / analysis through the ZymoBIOMICS pipeline (Zymo, Inc; Irvine CA). Amplicon sequence varients were generated via DADA2 and aligned to the Zymo Research Database for taxonomy assignment. QIIME2 (2023.9.2) was used for composition assessment, alpha-diversity, and beta-diversity analyses, Uclut with Zymo Research Database was used for taxonomy assignment. All data was visualized using GraphPad PRISM software. Demultiplexed FASTQ files are available in the NIH SRA database. Complete vendor analyzed datasets are available through Zenodo.

### 3-phenoxybenzoic acid (3-PBA) quantification

#### Serum

Serum collected at the indicated endpoints was assessed using on-line solid phase extraction (on-line SPE) liquid chromatography (LC) coupled with negative mode electrospray ionization (ESI)-tandem mass spectrometry (MS/MS) with isotope dilution quantification. Sample preparation involves diluting serum 50-fold with Milli-Q water, followed by spiking 200µL of the diluted sample with a stable isotopic analogue of the target analyte. Next, 300μL of 1,500 units/mL b-glucuronidase/sulfatase in 0.2M sodium acetate buffer, pH 5 is added to all samples. Samples are capped, vortex mixed, and placed in a 37ᵒC incubator, rotating at 150 rpm, overnight (∼15hrs), to liberate the target analyte from its conjugated form. After incubation, the enzymatically digested samples are centrifuged, transferred to autosampler vials, and injected onto a column switching system for extraction of the target analyte on a Strata RP on-line solid phase extraction (SPE) column (2.1 x 20mm). The on-line SPE column is washed with a mixture of acetonitrile: Milli-Q water (10:90, v/v) solution to remove undesired matrix interferences. Then, the target analyte is eluted from the on-line SPE column to a Poroshell 120 EC-C18 analytical column (3.0 x 100mm, 2.7 um) for chromatographic separation. The mobile phase consists of A) 0.1% acetic acid B) acetonitrile.

#### Urine

Urine was assessed using on-line SPE-LC coupled with negative mode ESI MS/MS) with isotope dilution quantification. Sample preparation involves diluting urine 500-fold with Milli-Q water, followed by spiking 200µL of the diluted sample with a stable isotopic analogue of the target analyte. Next, 300μL of 1,500 units/mL b-glucuronidase/sulfatase in 0.2M sodium acetate buffer, pH 5 is added to all samples. Samples are capped, vortex mixed, and placed in a 37ᵒC incubator, rotating at 150 rpm, overnight (∼15hrs), to liberate the target analyte from its conjugated form. After incubation, the enzymatically digested samples are centrifuged, transferred to autosampler vials, and injected onto a column switching system for extraction of the target analyte on a Strata RP on-line solid phase extraction (SPE) column (2.1 x 20mm). The on-line SPE column is washed with a mixture of acetonitrile: Milli-Q water (10:90, v/v) solution to remove undesired matrix interferences. Then, the target analyte is eluted from the on-line SPE column to a Poroshell 120 EC-C18 analytical column (3.0 x 100mm, 2.7um) for chromatographic separation. The mobile phase consists of A) 0.1% acetic acid B) acetonitrile.

### Deltamethrin quantification via GC-HRMS analysis

*Ex vivo* cultures treated with 1mM deltamethrin (as described above) were sampled at the indicated timepoints. Samples were thawed at 4°C, and 100μL culture aliquots were treated with 25μL formic acid and extracted by adding 200μL (4:1 hexane/ethyl acetate) containing ^13^C labeled internal standards (ISs). Treated samples were vortexed for 1hr, and then centrifuged for 10min at 4000rpm, at 4°C. After centrifuging, 150μL of supernatant was transferred to a new 96-well plate containing ∼20 mg of MgSO4, PSA, and C-18, vortexed for 2 min, and centrifuged for 10min at 4000rpm. The resulting supernatant was transferred to a 96-well plate containing glass inserts, and analyzed immediately.

To detect deltamethrin, untargeted analysis was performed using a Trace 1610 gas chromatography system coupled to an Exploris GC-HRMS system (Thermo Fisher Scientific, Rockford, IL, USA). Separation was accomplished by injecting 6µL of extract into a PTV inlet with a variable temperature program connected to a 15m DB-5MS column equipped with a 10m guard column (Restek), 1.2 mL/min flow rate, and the following oven temperature gradient: 50°C hold for 2.5min, increase to 320°C at 25°C/min and hold for 4min. The HRMS resolution was set at 60,000 FWHM (at m/z 200). The scan range was set from 50 to 75 m/z; the automatic gain control (AGC) target and maximum injection time in full-scan MS settings were set to Standard and Auto, respectively. This allowed the instrument to automatically adjustment the injection time for an ion count of ∼1 × 10^6^ for MS. The electron ionization source was maintained at 280°C and 70 eV during the course of the run. The mass spectrometer was calibrated weekly, the source was cleaned weekly, and columns were replaced weekly or every three batches. The peak for deltamethrin was integrated using TraceFinder (Thermo Fisher Scientific) and identified by comparison to an authentic reference standard using a mass and retention time error threshold of 5 ppm and 20 seconds, respectively.

### Hepatic bulk RNAseq

RNA was extracted from isolated livers (left medial lobule), collected at the indicated timepoints, using the Qiagen RNeasy Kit (#74104) according to manufacturer’s guidelines. Untargeted cDNA libraries were generated using Takara SMART-Seq mRNA HT LP kits (#634792) per manufacturer’s protocol. to generate bulk RNA-sequencing libraries for each sample. Libraries were sequenced by Admera Health (South Plainfield, NJ) on a NovaSeq X Plus to a sequencing depth of 50M reads per sample, with 150bp paired end reads.

Quality control of sequencing was performed on data from Admera using FastQC (version 0.11.9). Post QC reads were pseudo-aligned using Kallisto (version 0.44.0) to a mouse reference genome (GRCm39) using default parameters. After alignment, files were imported into R (version 4.4.3) and the analysis workflow of DIY Transcriptomics followed (Berry et al., 2021). Briefly, gene counts were converted into TPM normalized log2 counts per million, and genes expressed in less than two samples were excluded. Differential gene expression analysis was performed using limma (version 3.62.1) and edgeR (version 4.4.2). Genes were considered to be differentially expressed in each comparison if they had a Bonferroni-adjusted *p*-value less than 0.05 and a log2 fold change ≥|1.0|. Pathway analysis was performed using Metascape (Zhou et al., 2019). Hepatic transcriptomics are available in the NIH SRA database; all DEG and pathway analysis available in the supplemental tables associated with this manuscript and on Zenodo.

### Colonic bulk RNAseq

RNA was extracted from ∼2cm of the proximal colon via TRIzol-chloroform extraction. Tissue was sonicated in 400μL TRIzol (Qiagen, Cat#: 79306) on ice. 100µl chloroform was added, and samples were gently mixed via inversion. Following a 3min incubation at RT, samples were centrifuged at 12k rpm for 15min at 4^°^C. The upper aqueous phase was transferred to a Qiashredder tube (Qiagen, Cat#: 79656) and centrifuged for 2min at 16k rpm. Following extraction, samples were processed using the RNeasy Mini Kit (Qiagen, Cat#: 74104) per supplier’s protocol. Samples were eluted with 40μL RNase-free water and provided to Novogene Corporation, Inc (Sacramento, CA) for library preparation, sequencing and analysis. Libraries were prepared with the NEBNext UltraTM RNA Library Prep Kit for Illumina, with poly-T enrichment and sequenced on a NovaSeq X Plus to a depth of ∼50M reads per sample with 150bp paired-end reads. Colonic transcriptomics are available in the NIH SRA database; all DEG and pathway analysis available in the supplemental tables associated with this manuscript and further protocols and analysis files on Zenodo.

### Quantitative Real-Time Polymerase Chain Reaction Analysis (qRT-PCR)

*Ces2a* expression was measured in a targeted fashion using qRT-PCR. RNA was extracted via TRIzol-chloroform extraction with further purification using an RNeasy Mini Kit (Qiagen, Cat#: 74104), as described above. cDNA was synthesized using Applied Biosystems™ High Capacity cDNA kit (Applied Biosystems, Warrington, UK) according to the manufacturer’s instructions and assessed via qRT-PCR using SYBR® Green PCR Master Mix (Applied Biosystems), using the primers in the Key Resource Table for *Ces2a* with *Gapdh* as the control on a 7900HT Fast Real-Time PCR System. Expression level was calculated by transforming the cycle threshold value (Ct) using the 2−ΔΔCT method.

## QUANTIFICATION AND STATISTICAL ANALYSIS

### Statistical Analysis

Data are presented as mean ± SEM, unless indicated otherwise within the figure legend. Individual points represent biological replicates from an individual mouse tissue or stool. Statistical tests were performed using GraphPad Prism 10 statistical software (RRID: SCR_002798; v10.4.1). As indicated in figure legends, non-transcriptomic comparisons were made using two-way ANOVA (treatment, time; or treatment, colonization status) and appropriate post-hoc test, or for mono-colonization qPCR study, a one-way ANOVA with post-hoc tests, as indicated in figure legends. Bulk transcriptomics were assessed as described above with DEGS assigned following FDR controlled *p*-value adjustment using Benjamini-Hochberg correction within the DESeq2 R package (RRID: SCR_015687). All source data are available within the supplemental tables included with this manuscript, at Zenodo, or available on NIH SRA.

## Notes

### Competing Interest Statement

The authors have declared no competing interest.

