## Supplemental Figures S1 - S4 for "The gut microbiome promotes detoxification responses to an environmental toxicant"

#### Supplemental Information

**Supplemental Figure S1 *associated with Figure 1*.** Deltamethrin directly impacts a complex stool-derived microbiome and modulates growth *in vitro* of select taxa.

**Supplemental Figure S2, *associated with Figure 2*.** The native microbiome impacts hepatic gene expression.

**Supplemental Figure S3, Supplemental Fig. S3, *associated with Figure 2.*** CYP enzymes show variable responses across exposure and microbiome status

**Supplemental Figure S4, *associated with Figure 3*.** Microbiome modulation of colonic transcriptomes.

### Supplemental Figure S1

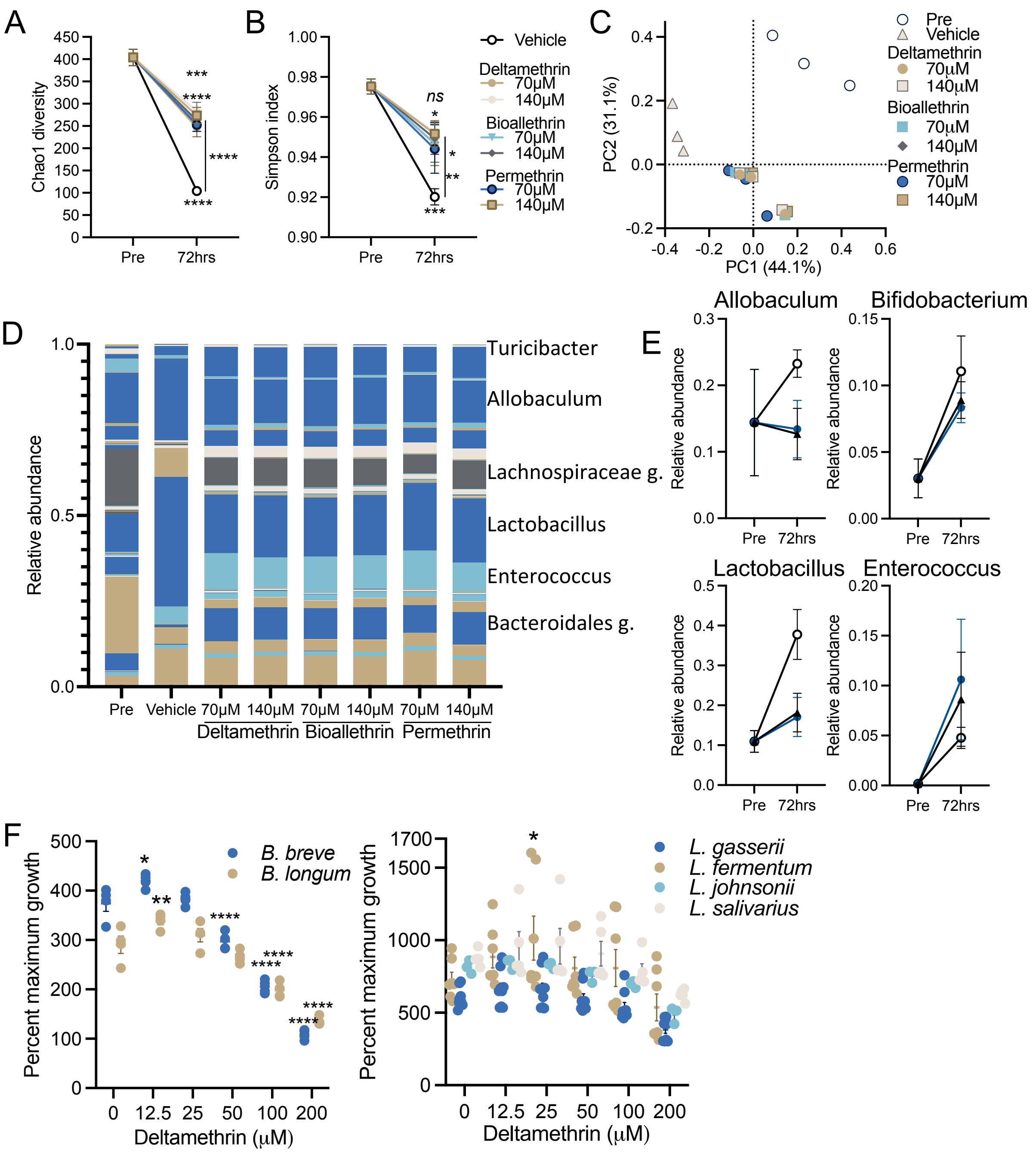


**Supplemental Figure S1, *associated with Figure 1*. Deltamethrin directly impacts a complex stool-derived microbiome and modulates growth *in vitro* of select taxa.** Stool derived from wildtype mice was cultured anaerobically with 70µM or 100µM of deltamethrin, bioallethrin, or permethrin, and analyzed by 16S RNA sequencing prior to exposure or 72hrs post-exposure. Alpha diversity measurements for Chao1 (**A**) and Simpson indexes (**B**). (**C**) PCA plots of Bray-Curtis beta diversity measures. (**D**) Genera composition charts. (**E**) Select genera displaying time or treatment effects following deltamethrin exposure. (F) Single type strains of the indicated species were grown to their maximum optical density (OD600) in the presence of indicated deltamethrin concentrations (19hrs *Bifidobacteria,* 12hrs *Lactobacillus*). N=8 (A-E), N= 3-9 (F). A, B, E points represent mean and bars the standard error. **C, F** points represent independent samples. A, B data assessed by two-way, repeated measures ANOVA and post-hoc Fisher's tests, with lines indicating between group differences. F data assessed by two-way ANOVA with Sidek's posthoc test to each species untreated group *$p \leq 0.05$; **$p \leq 0.01$, *** $p \leq 0.001$; **** $p \leq 0.0001$

### Supplemental Figure S2

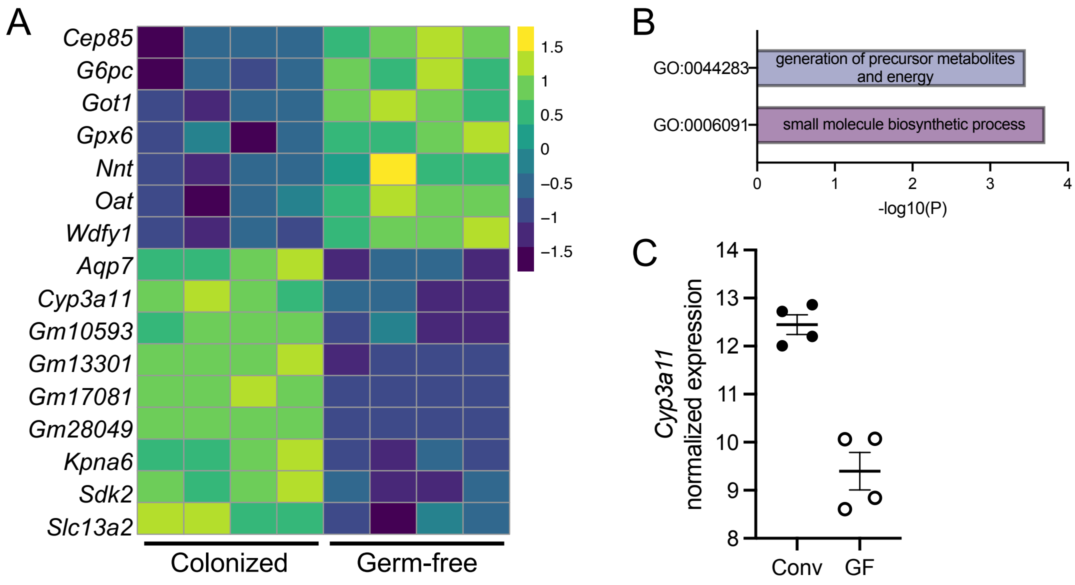


**Supplemental Figure S2, *associated with Figure 2*. The native microbiome impacts hepatic gene expression.** RNAseq analysis of liver tissue derived from conventional and germ-free mice. **A)** Heatmap of all DEGs meeting both adjusted *p*-value threshold and fold-change. **B)** Metascape pathway analysis of enriched GO terms of genes increased in germ-free liver tissue. **C)** Normalized expression of *Cyp3a11* isolated from (**A**). N=4, **A, C,** points represent individual mice and bars the mean and standard error. See Supplementary Table S2

### Supplemental Figure S3

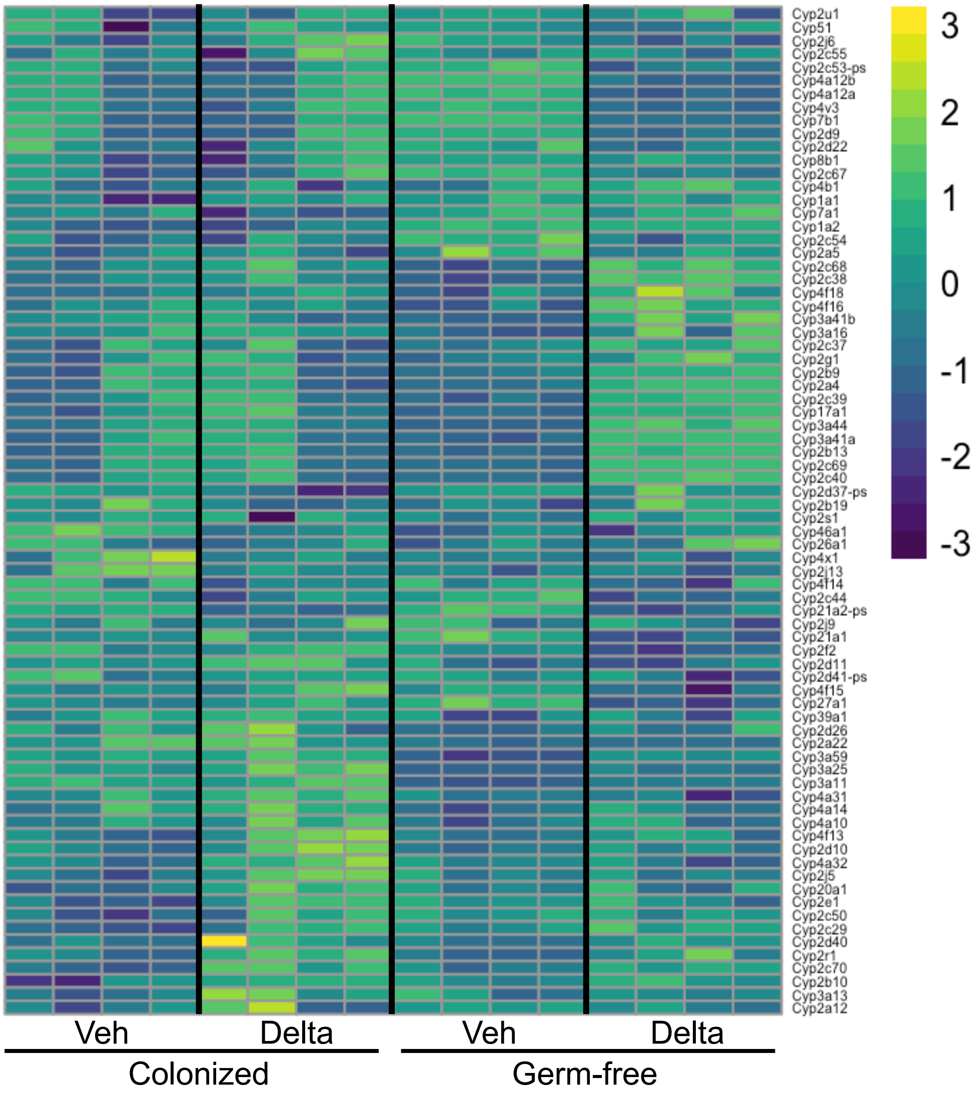


**Supplemental Fig. S3, *associated with Figure 2.* CYP enzymes show variable responses across exposure and microbiome status.** Expression level of detected genes for CYP enzymes in the liver of conventional and germfree mice. N=4.

### Supplemental Figure S4

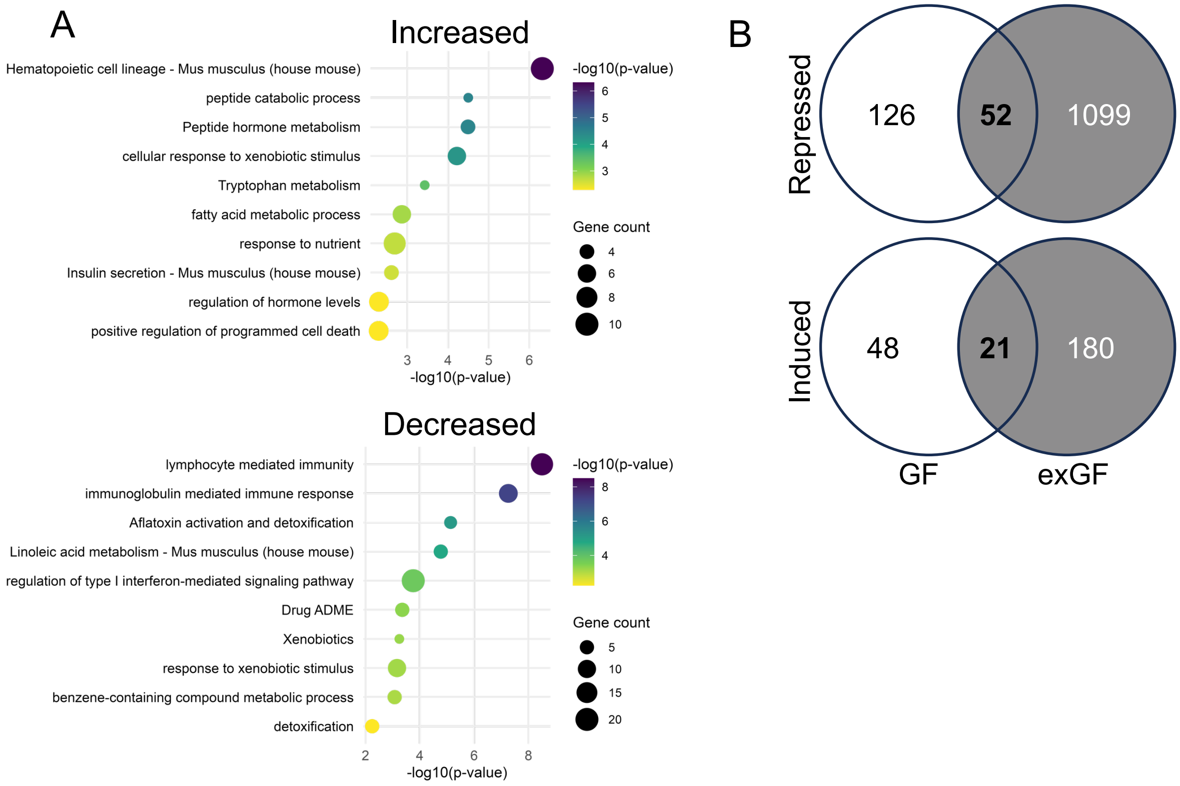


**Supplemental Fig. S4, *associated with Figure 3*. Microbiome modulation of colonic transcriptomes. A)** RNAseq analysis of colon tissue derived from conventional and germ-free mice, with Metascape pathway analysis of enriched terms in both increased and decreased genes in GF mice. **B)** Comparison of colonic transcriptomes between GF and exGF mice, as they differ from colonized controls.
